# Efficient and Economical Targeted Insertion in Plant Genomes via Protoplast Regeneration

**DOI:** 10.1101/2021.03.09.434087

**Authors:** Chen-Tran Hsu, Yu-Hsuan Yuan, Yao-Cheng Lin, Steven Lin, Qiao-Wei Cheng, Fu-Hui Wu, Jen Sheen, Ming-Che Shih, Choun-Sea Lin

## Abstract

Versatile genome editing can be facilitated by the insertion of DNA sequences into specific locations. Current protocols involving clustered regularly interspaced short palindromic repeats (CRISPR) and CRISPR-associated (Cas) proteins rely on low efficiency homology-directed repair or non-homologous end joining with modified double-stranded DNA oligonucleotides as donors. Our simple protocol eliminates the need for expensive equipment, chemical and enzymatic donor DNA modification or plasmid construction by using polyethylene glycol-calcium to deliver non-modified single-stranded DNA oligonucleotides and CRISPR-Cas9 ribonucleoprotein into protoplasts. Plants regenerated via edited protoplasts achieved targeted insertion frequencies of up to 50.0% in *Nicotiana benthamiana* and 13.6% in rapid cycling *Brassica oleracea* without antibiotic selection. Using a 60-nt donor containing 27 nt in each homologous arm, 6 of 22 regenerated *N. benthamiana* plants showed targeted insertions, and one contained a precise insertion of a 6-bp *Hind*III site. The inserted sequences were transmitted to the next generation and invite the possibility of future exploration of versatile genome editing by targeted DNA insertion in plants.

## Introduction

To insert DNA into a specific location in the plant genome, a DNA double-strand break (DSB) must be induced at the target position to facilitate the donor DNA (DD) insertion into this position via homology-directed repair (HDR)^1–12^ or non-homologous end joining (NHEJ).^13–15^ Many tools for creating DSBs are currently available.^16–18^ Due to its versatility and simplicity, the combination of clustered regularly interspaced short palindromic repeats (CRISPR) and CRISPR associated (Cas) proteins has become a favorite approach among genetic engineers.^4^ In addition to CRISPR-Cas reagents, increasing the amount of delivered DD enhances targeted insertion (TI) efficiency. For example, in maize (*Zea mays*), targeted mutagenesis using Cas9 and guide RNA (gRNA) has been achieved via different methods, although TI plants were obtained only via biolistic methods rather than *Agrobacterium*-mediated transformation because the number of DD copies delivered by the latter method was low.^2^

Protoplasts offer an alternative system for genome editing and genetic transformation, since they enable the delivery of high numbers of DD copies^19, 20^ to enhance TI in the genome before plant regeneration.^19^ Like the transcription activator-like effector nuclease genome editing system^21^, CRISPR reagents can be delivered into protoplasts via polyethylene glycol (PEG)-Ca^2+^-mediated transfection of ribonucleoprotein (RNP) or plasmid DNA, and targeted mutations can be achieved.^22–29^ The mutated protoplasts can then be regenerated into plants without chimerism and the mutated alleles are passed onto the progeny.^22, 24, 25^ Here, we describe a simple, high-efficiency TI method based on the protoplast strategy for genome editing in plants using Ca9-gRNA and non-modified synthetic single-stranded oligonucleotide DNA (ssODN) DDs that does not require expensive equipment.

## Materials and Methods

### Protoplast isolation, transfection, and regeneration

*Nicotiana benthamiana* and rapid cycling *Brassica oleracea* (RCBO) plants were propagated in ½ -strength Murashige and Skoog (½ MS) medium supplemented with 30 g/L sucrose and 1% agar under a 12-h/12-h light/dark cycle at 25°C. Protoplast isolation was performed according to Lin *et al.*^24^ and Hsu *et al.* ^29^ except for the digestion solution [½ MS medium supplemented with 1 mg/L 1-naphthaleneacetic acid (NAA), 0.3 mg/L kinetin, 30 g/L sucrose, 0.4 M mannitol (1N0.3K), 1% cellulose, 0.5% Macerozyme] and digestion time (3 days) in *N. benthamiana*. The protoplasts were co-transfected with RNP and 50 μg synthetic single-stranded oligonucleotide DNA (ssODN; Genomics, Taipei, Taiwan) according to Woo *et al.*^22^ Transfected protoplasts were incubated in a 5-cm-diameter Petri dish containing liquid callus medium (1N0.3K) for 3 weeks. RCBO calli were additionally incubated in 1 mg/L NAA, 1 mg/L 6-benzyladenine (BA), and 0.25 mg/L 2,4-dichlorophenoxyacetic acid for 3 days in the dark. The calli were transferred to liquid shooting medium (containing 2 mg/L BA for *N. benthamiana*, 0.1 mg/L thidiazuron for RCBO) in a 9-cm-diameter Petri dish and incubated at 25°C for 3–4 weeks in the light (16-h/8-h light/dark, 3000 lux). Green explants larger than 5 mm were incubated in solid shooting medium and sub-cultured every 4 weeks. Shoot clusters with leaves were then transferred to solidified rooting medium (HB1: 3 g/L Hyponex No. 1, 2 g/L tryptone, 20 g/L sucrose, 1 g/L activated charcoal, 10 g/L agar, pH 5.2).

### Cas9 protein purification, single-guide RNA synthesis, and Cas9 RNP nucleofection

Preparation of Cas9 protein and single-guide RNA (sgRNA) and Cas9 RNP nucleofection were performed according to Huang *et al.*^30^ Cas9 recombinant protein was overexpressed in *Escherichia coli* BL21 harboring the plasmid pMJ915 (Addgene # 69090). Cas9 protein was purified and stored at −80°C in Cas9 RNP buffer (20 mM HEPES at pH 7.5, 150 mM KCl, 10% glycerol, and 1 mM β-mercaptoethanol). The sgRNAs were synthesized by *in vitro* transcription (IVT) using T7 RNA polymerase (New England Biolabs; M0251L). The DNA oligonucleotides used for IVT template assembly are listed in Table S1. The final sgRNA products were dissolved in Cas9 RNP buffer, quantified using a NanoDrop Lite (Thermo Fischer Scientific), and stored as aliquots at −80°C. Cas9 RNP complexes were assembled immediately before nucleofection by mixing equal volumes of 40 μM Cas9 protein and 88.3 μM sgRNA at a molar ratio of 1:2.2 and incubating at 37°C for 10 min.

### Validation of targeted insertions in protoplasts and regenerated plants

Genomic DNA was extracted from pooled protoplasts and regenerated plants using a Geno Plus Mini Genomic DNA Extraction Kit (GG2002, Viogene, New Taipei City, Taiwan). To amplify the genomic region targeted by the sgRNA, the corresponding pairs of primers were designed. Primer sequences are shown in Table S1. The PCR conditions were 94°C for 5 min, 35 cycles of 94°C for 30 s, annealing at 55–63°C for 30 s, polymerization at 72°C for 30 s, followed by 72°C for 3 min. The polymerase chain reaction (PCR) products were digested using the appropriate restriction enzyme or RNP and subjected to electrophoresis. The PCR products that could not be digested by restriction enzymes at target sites near the protospacer adjacent motif (PAM) sequence (*Bst*NI in *NbPDS1* E target site) or RNP were defined as edited. The PCR products that could be digested by the restriction enzyme in the DD and for which sequencing was confirmed are defined as TI. The PCR products were cloned into the T&A vector (FYC002-20P; Yeastern Biotech, New Taipei City, Taiwan). Putative colonies containing the edited DNA were confirmed by Sanger sequencing.

### Whole-genome sequencing for off-target DD insertion analysis

Leaves of *N. benthamiana* protoplast regenerated plants were collected for genomic DNA purification. Genomic DNA for genome sequencing was extracted using a Plant DNA Purification Kit (DP320, Tiangen, https://en.tiangen.com). Paired-end libraries of DNA were constructed by the NEBNext Ultra II DNA Library Prep for Illumina Kit (New England Biolabs; E7645L) with 2 × 150 bp with average insert size ~900 bp and sequenced on a NovaSeq 6000 platform (Illumina; 20028312). Three technical replicates were performed for each sample. Total reads were 120 Gbp per regenerated plant and the sequencing depth was more than 30×. To ensure the read quality, the first 10 bases of Illumina reads were removed and the last 141 bases were retained for further analysis. High-quality Illumina reads were aligned with the *N. benthamiana* genome (genome assembly v.1.0.1) by BWA (v.0.7.17) with default setting. SNPs and INDELs were identified by DeepVariant (v.1.1.0-GPU, WGS model) and subsequently processed by GLnexus (v.1.2.7, DeepVariantWGS model) and bcftools (v.1.10.2, FMT/GQ<=20 and GT= “RA”) for a joint variant calling. Off-target sites were predicted by Cas-OFFinder (v.2.4.1) with default settings. Sample specific SNPs and INDELs were compared with predicted off-target sites from Cas-OFFinder to identify coincident sites. To identify the potential off-target insertion sites, target sequence (Exp. 1 DD sequence, TTTGCGATGCCTAACAAGCTTCAGGGGGAGTTCAGCCGCTT) was used as the query in a high sensitivity BLASTN search strategy (-dust no -soft_masking false -word_size 4 -gapopen 1 -gapextend 2 -penalty -1 -reward 1 -evalue 5000 - perc_identity 80 -num_alignments 50000) against the Illumina reads. The high-scoring segment pairs of Illumina reads were filtered according to the known edited sequence lengths in different samples. Candidate Illumina reads were then retrieved to further examine the exact location in the *N. benthamiana* genome by BLASTN (-dust no -soft masking false -task blastn-short -evalue 0.1 -perc_identity 90 - num_descriptions 1 -num_alignments 1). If the Illumina read was identical to the published genome sequence, it was concluded that these sequences were the same as the DD already existed and were not caused by TI. If there was a difference from the published WT genome sequence, and the difference was the same as the DD, it was regarded as an off-targeted insertion. The raw reads were deposited in the NCBI SRA database (BioProject: PRJNA667297; https://www.ncbi.nlm.nih.gov/bioproject/PRJNA667297).

## Results and Discussion

The protoplast regeneration protocol (Figure 1) used in the present study was modified from previously published protocols.^24, 25, 29^ A key step to our approach is the observation that the phase of the cell cycle largely governs the choice of pathway used for DNA repair: NHEJ is the major DNA repair pathway during the G1, S, and G2 phases, whereas HDR occurs only during the late S and G2 phases^31, 32^. Cell cycle synchronization is an effective strategy for enhancing TI efficiency in human embryonic kidney 293T cells.^33^ Here, 5-ethynyl-2′-deoxyuridine (EdU) staining was used for detection of S-phase cell-cycle progression. To increase the number of cells in the late S and G2 phases, *N. benthamiana* leaves were incubated in 1N0.3K solid medium for 3 days before protoplast isolation (Figure S1A). In comparison with 1N0.3K treatments, no EdU signal was identified in protoplasts incubated in ½ MS, 0.4 M mannitol solid medium (Figure S1B).^34^ Based on single cell analysis^24^, the TI efficiency increased after incubation in 1N0.3K (Figure S1C). For *N. benthamiana*, to simplify the procedure, we used 1N0.3K for digestion solution preparation and incubated the cut leaf material for 3 days.

**Figure 1.**
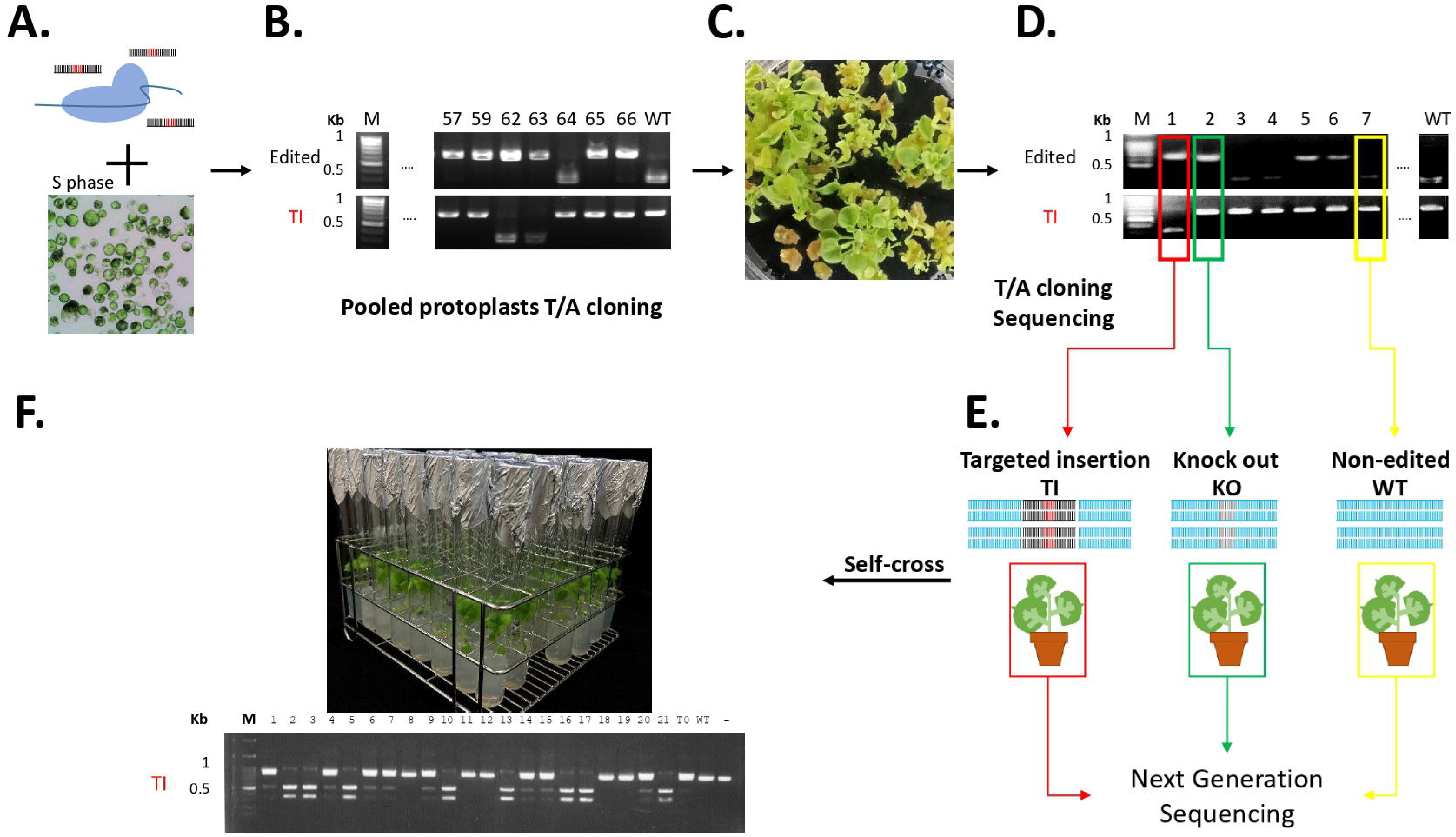
Strategy for targeted DNA insertion using ribonucleoprotein and single-stranded oligonucleotide DNA in *Nicotiana benthamiana*. (A) Ribonucleoprotein (RNP) and single-stranded oligonucleotide DNA (ssODN) are delivered to S-phase protoplasts by polyethylene glycol-Ca^2+^–mediated transfection (Figure S1). (B) Pooled transfected protoplasts DNA was isolated and the target gene was amplified by polymerase chain reaction (PCR) and cloned into the T/A vector for genotyped using restriction enzymes or RNP. (C) Transfected protoplasts are regenerated. (D) DNA from the regenerated plants is amplified by the polymerase chain reaction and genotyped using restriction enzymes or RNP. (E) DNA is purified from targeted insertion (red), knockout (green), and non-edited (yellow) regenerated plants and sequenced to determine whether ssODN was inserted into other positions. (F) Offspring are genotyped and sequenced to test whether the inserted sequence is heritable.

To bypass the need for plasmid construction, we used RNP as the Cas9-gRNA reagent. For the DD, we used short non-modified synthetic ssODN, which is relatively inexpensive and easy to obtain. The RNP and ssODN were delivered into protoplasts using PEG-Ca^2+^-mediated transfection (Figure 1A). ^20, 22^ After 3 days of incubation in 1N0.3K liquid callus medium, genomic DNA was isolated from the protoplasts and the target gene was amplified by PCR and cloned into the T/A vector for genotyping and Sanger sequencing to assess the TI efficiency (Figure 1B). These protoplasts were cultured and regenerated (Figure 1C).^24, 25, 29^ The rooted plants were incubated in the growth chamber and genotyped (Figure 1D). These regenerated plants grew normally and produced seeds. DNA from two types of edited and regenerated plants, carrying TI or knockout (KO), and one non-edited (WT) and regenerated plant were purified for genome-wide sequencing to assess the presence or absence of off-target DD insertion as well as genome stability through the protoplast regeneration processes (Figure 1E). DNA from the TI T_1_ progeny was extracted for genotyping to test whether the inserted fragment was heritable (Figure 1F).

For the regenerated plants with TI, we conducted experiments using *PHYTOENE DESATURASE1* (*NbPDS1*) as a target site in *N. benthamiana* (Figure S2).^35^ Based on single cell analysis^24^, the TI only occurred when protoplasts were transfected with DD and RNP (Figure S2A). To evaluate the effect of the length of the homologous arms and the total length of ssODN on TI efficiency at the expected sgRNA target position, we synthesized the DD of 20 nt, 40 nt, or 60 nt ssODN carrying a *Hind*III site and added 7, 17, or 27 homologous arms, respectively, on the left and right sides (Figure S2B). We genotyped *NbPDS1* PCR products from the edited and regenerated plants. No TI were identified with the 20-nt ssODN in regenerated plants, whereas the 40-nt and 60-nt ssODNs produced TI in regenerated plants with efficiencies of 27.3–31.8% without antibiotic or phenotypic selection (Figure S2C, D). These results suggested that the length of the ssODN donor is an important factor in determining the TI efficiency. In a previous study, a 59-nt ssODN (ssADHE) failed to give rise to successful insertions in 23 T_0_ rice (*Oryza sativa*) plants.^13^ There were no homologous arms in ssADHE and a lower DD concentration was used than was applied to the protoplasts during RNP transfection in the current study. Based on these results, we selected 40 nt, which had highest TI efficiency, as the length of the ssODN in our subsequent experiments, except for Experiment (Exp.) 2 (44 nt). Interestingly, one of the regenerated *N. benthamiana* TI lines mediated by the 60-nt ssODN had a precise insertion of 6 bp *Hind*III sequence (8.3%) in the predicted target site (+27#6, Figure S2E, F).

Next, we examined the effect of the insertion length in a 40-nt DD on TI efficiency (Figure 2). In Exp. 1, the *Hind*III site was generated with additional 2-nt insertion in the DD (Figure 2A), and TI efficiency in the regenerated plants was 18.2% (Figure 2B, C). In Exp. 2, the PAM sequence in DD was mutated, 6 nt were added, and the TI efficiency increased to 50.0%. We also increased the insertion length to 15 nt from 6 nt in Exp. 3, including sites for *Nhe*I and *Bam*HI endonucleases in the insertion, which enabled us to confirm the integrity of TI genotyping by restriction enzyme digestion. As the insertion length increased and the length of the homologous arms decreased (11 bp and 14 bp, respectively), the TI efficiency decreased slightly (40.9%). Three regenerated plants contained only the *Nhe*I site, indicating partial DD insertion (Figure 2B). Perhaps the ssODN DD was unstable in protoplasts and had been partially degraded before insertion, which would cause the inserted sequence to be incomplete. There was no phosphorothioate-linkage modification in the DD we used in this study, which may have led to the partial degradation of DD before or during the TI process.^13^

**Figure 2.**
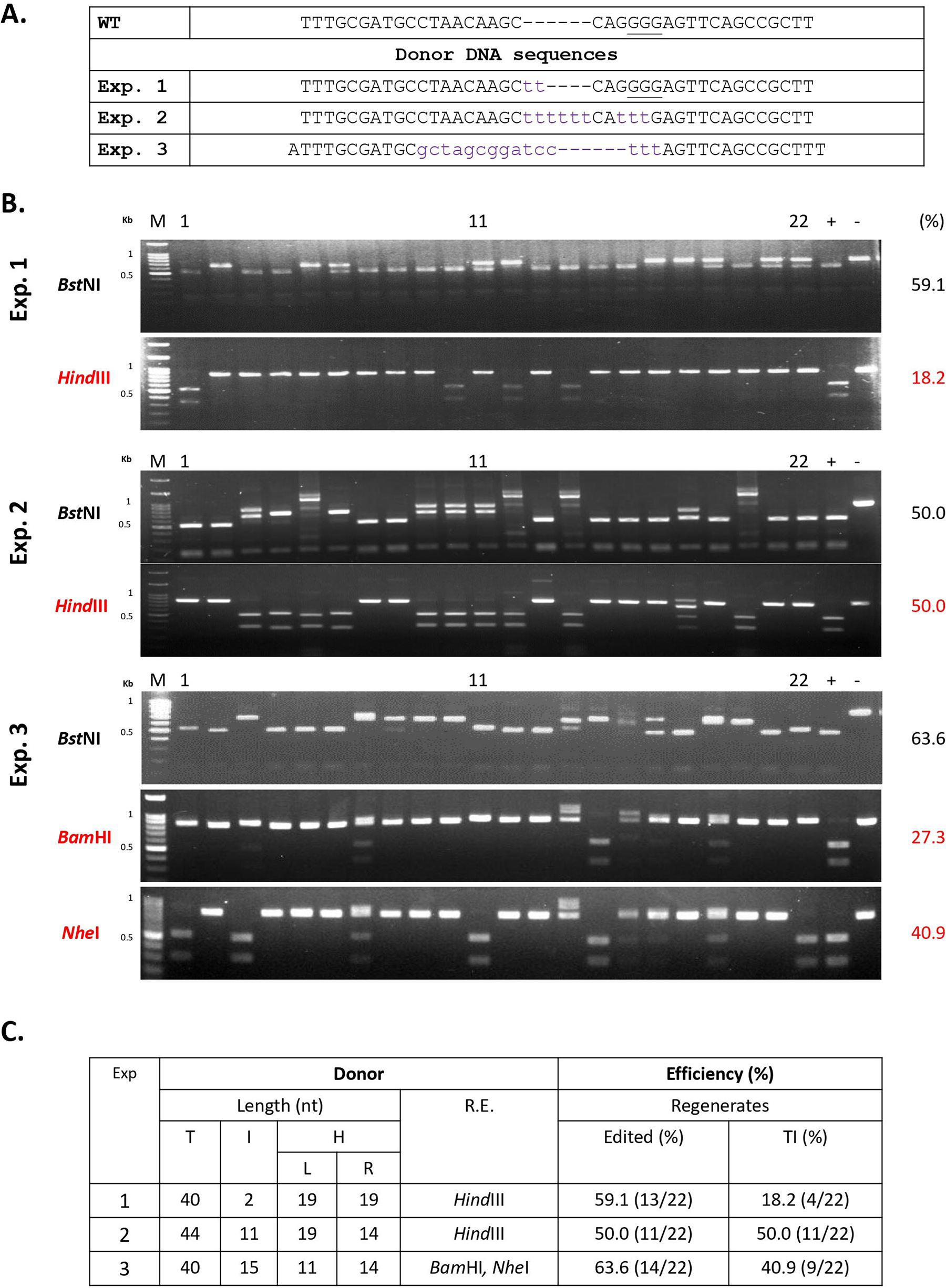
Targeted insertion in *NbPDS1* facilitated by short ssODN with different sequences and homology arm length in regenerated *Nicotiana benthamiana* plants. (A) Donor sequences used in Exp. 1–3. Lowercase: insertion or replacement nucleotides. The protospacer adjacent motif (PAM) is underlined. (B) Polymerase chain reaction (PCR) of the target gene in the regenerated plants was performed in each experiment, and different restriction enzymes were used for genotyping. (black: *Bst*NI, edited; red: *Hind*III, targeted insertion). M: marker. -: wild-type control. +: restriction enzyme control. Targeted insertion PCR product sequences are shown in Tables S2–S4. Because the insertion DNA could be cleaved to ~20 bp by restriction enzymes which we designed in DD, all of the TI regenerated plants had the same digested band pattern. In most of cases, the *Bst*NI site in the target site is disrupted by targeted mutagenesis and TI. However, if cytosine is the last nucleobase of TI, for example: Exp. 1#1, 10, 14, the *Bst*NI site is retained (CCAGG) and PCR products could be cleaved. (C) Edited and targeted insertion efficiencies of different ssODN donors. T: total length of DD. I: insertion. H: homologous arm. L: left arm. R. right arm. R. E.: restriction enzyme.

We sequenced all of the *NbPDS1* genes with TI in the regenerated plant produced in Exp. 1, 2, and 3 (Tables S2, S3, and S4). In a few regenerated plants with TIs, one end matched the ODN precisely whereas the other end did not (Exp. 1#12 and Exp. 2#18). In these regenerated plants, the insertion size was 29–445 bp. These differences were caused by insertion of 1–13 repeats of the ssODN, although some of the repeats were incomplete. In rice, only 20% of TI plants had repeat insertions when using modified double-stranded DD.^13^ Both orientations were observed for the TI in our regenerated plants. Forward and reverse insertions have also been identified in rice.^13^

Accurate insertion or replacement of DNA fragments is vital for gene editing. Co-expressing two sgRNAs can lead to fragment deletion in protoplasts.^14, 24^ Therefore, two sgRNAs can be designed at both ends of an exon for exon replacement.^14^ To aid in TI and DNA replacement, we designed two sgRNAs (L1 and L2) based on both sides of the complementary strands of the original target site (E). These sgRNAs could form a combination of RNPs, including tail-to-head (L1+L2), tail-to-tail (L1+E), or head-to-head (E+L2) orientation (Figure 3A). The ssODN DDs were all located in the same strand of target site E (Figure 3B). In L1+L2 and E+L2, the sequence of ssODN contained 17 bp complementary to L2 sgRNA, which caused the L2 RNP to show reduced efficiency for generating DSBs (Figure S3). The notion that the strand of DD complementary to the target site can reduce RNP activity was validated by experiments using L1 and L2 with complementary ssODN. These results are different from those reported for human cell lines.^33^ In human cells, both strands of DD can be used for gene editing. The L1+E experiment had a higher fragment deletion rate than the others (Figure 3C). Except for a decrease in E+L2, the overall TI efficiency was similar to that using a single RNP (Figure 2C, Exp. 2 and 3).

**Figure 3.**
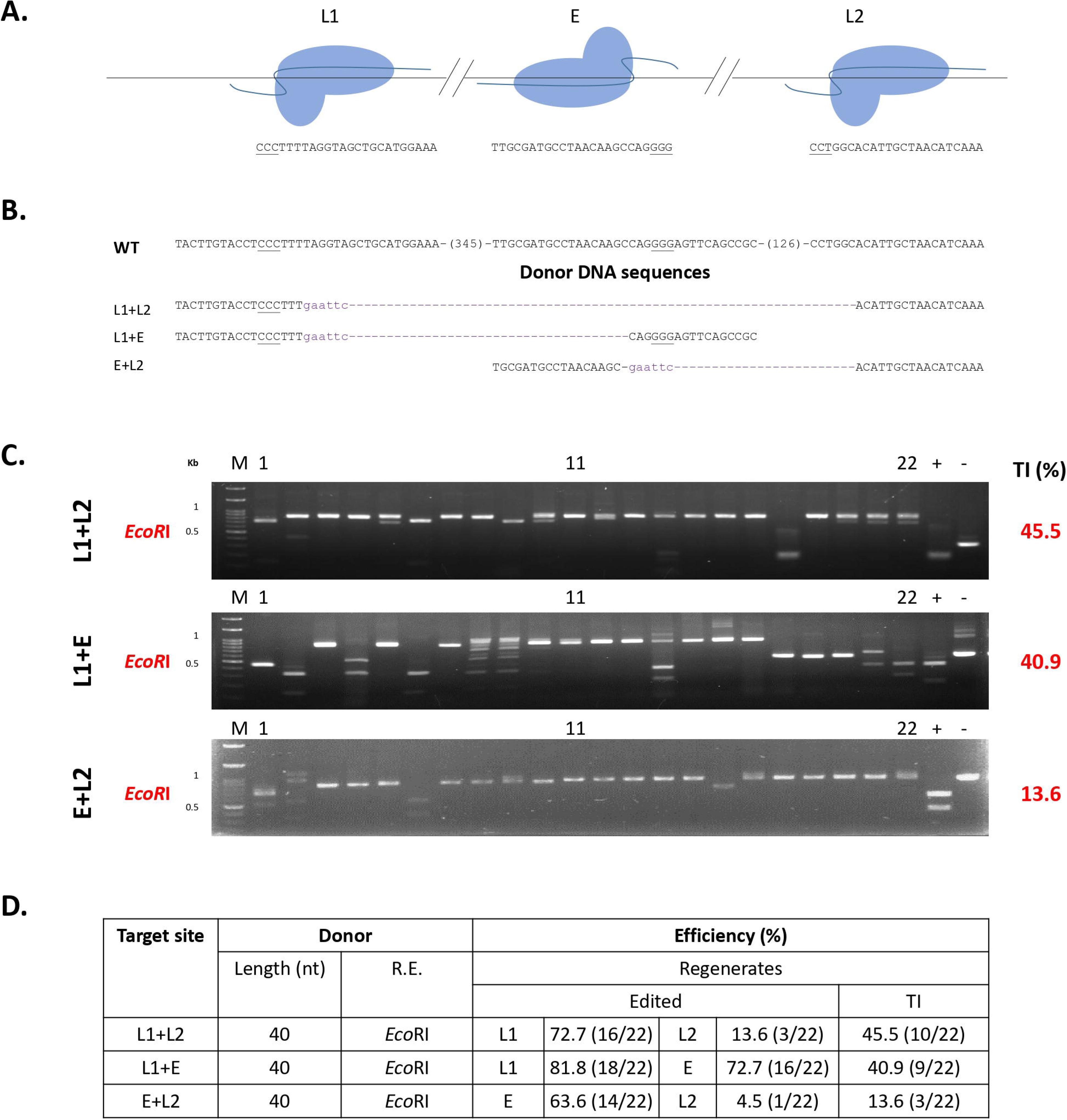
Targeted insertion using two ribonucleoproteins in *Nicotiana benthamiana*. (A) Relative positions and directions of the three ribonucleoproteins (RNPs) used. The target site is shown below. The protospacer adjacent motif is underlined. (B) Donor DNA (DD) used in each combination. Lowercase: inserted *Eco*RI site. (C) Restriction enzyme analysis of the target efficiency of regenerated plants derived from different RNP combinations and DD transfected protoplasts. TI: targeted insertion. (D) Summary of edited and TI efficiencies of RNP and single-stranded oligonucleotide DNA. R.E.: restriction enzyme.

Compared with the use of E RNP only, there was a decline in the TI/Edited ratio when two sgRNA RNPs were co-transfected into protoplasts (Figure 3D). The insertion sequences are shown in Tables S5, S6, and S7. From the results, it can be seen that by using 2 RNPs, a region of DNA can be successfully removed and inserted with DD. Therefore, this method is confirmed as applicable to exon replacement.^14^

To determine whether these insertions were heritable, we analyzed the progeny of five *N. benthamiana* regenerated plants (Figure S4A). We genotyped TI regenerated plant T_1_ seedlings and determined that all TI alleles were inherited (Figure S4B, C). Thus, these protoplast regenerated plants were not chimeric at the target gene. By contrast, in rice, most T0 plants appeared to be chimeric.^13^ In the current study, no *Cas9* gene was present in the genomes of the regenerated plants because we used RNP as the Cas9-gRNA reagent and therefore no new edited alleles were generated.

We also examined the TI efficiency in RCBO, targeting *BoSnRK1* and *BoGA4.a*^36^ (Figure 4). RCBO contains two *BoSnRK1* genes: *BoSnRK1.a* and *BoSnRK1.b* (Figure 4A). DD was inserted into the target sites (Figure 4B, C, Tables S8 and S9). Sequencing indicated that DD was inserted at an efficiency of 4.5–13.6% (Figure 4C), which is lower than that demonstrated in *N. benthamiana*.

**Figure 4.**
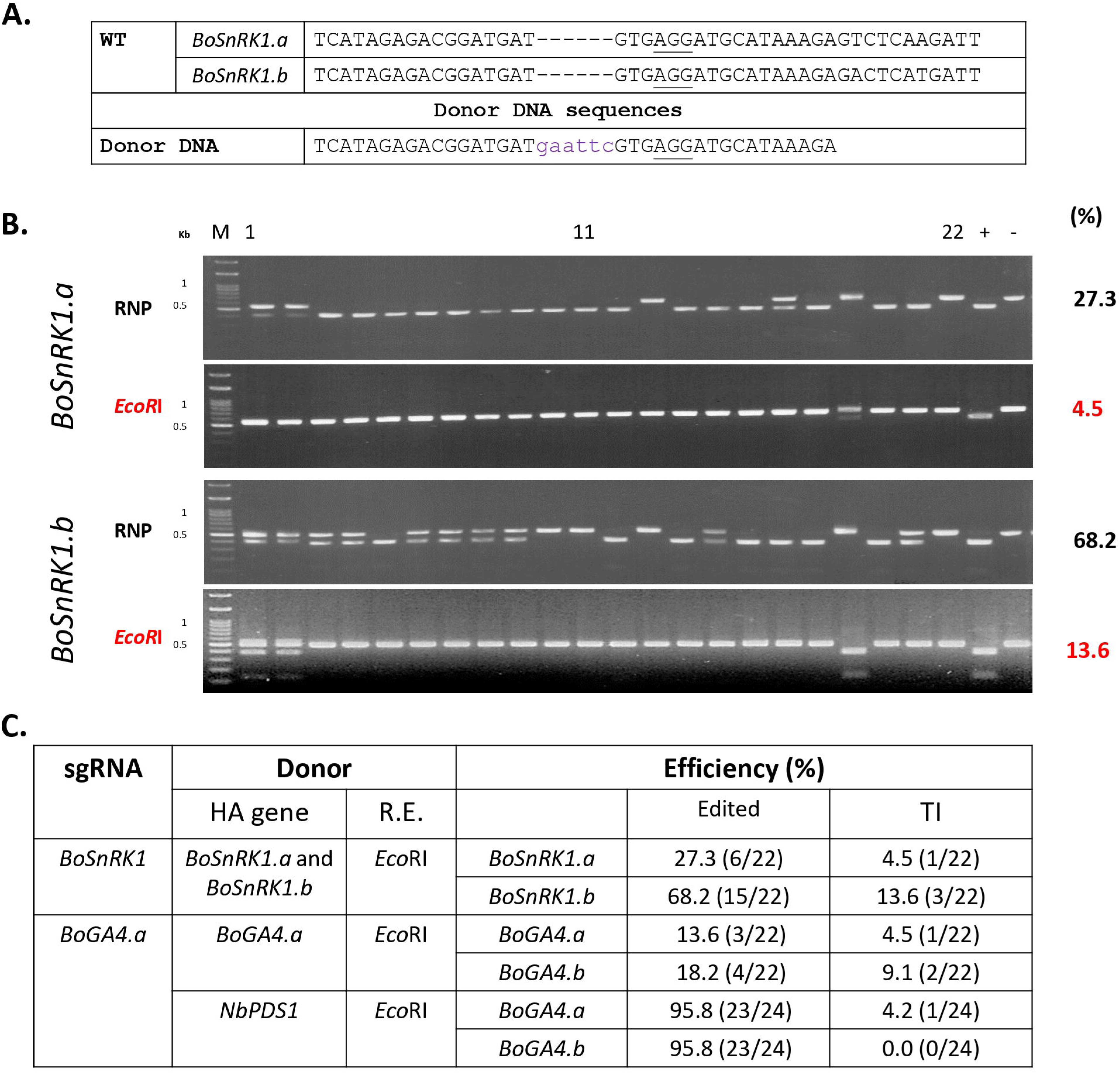
Targeted insertion using single-stranded oligonucleotide DNA donor DNA in rapid cycling *Brassica oleracea*. (A) Target site of *BoSnRK1* and the donor sequence. The protospacer adjacent motif is underlined. Lowercase: insertion sequence. (B) Genotyping of the regenerated plants derived from ribonucleoprotein (RNP) and single-stranded oligonucleotide DNA (ssODN) transfected protoplasts. The edited efficiency by RNP was assessed. Targeted insertion was assessed using the *Eco*RI site inserted by the ssODN. (C) Summary of edited and targeted insertion efficiencies. R.E.: restriction enzyme site inserted by ssODN. HA: homologous arm. TI: targeted insertion.

According to the results, we reasoned that the presence of a homologous arm on the DD designed for TI might not be necessary. To investigate whether ssODN DD with a non-homologous arm could be used for TI, we co-transfected RCBO protoplasts with *BoGA4.a* sgRNA RNP and *NbPDS1* DD. TI was observed in *BoGA4.a* in only 4.2% of the 24 regenerated plants (1/24), and none was observed in *BoGA4.b*, even though the editing efficiency was 95.8% (Figure 4C, Table S10). NHEJ is typically guided by short homologous DNA sequences (microhomologies) which affect joining efficiency greatly.^37^ The lower efficiency when using *NbPDS1* DD in RCBO in comparison with *N. benthemiana* is likely due to variations within genomes, target genes, or the microhomologies between DD and target site. It would require further investigation to understand the exact mechanisms.

*N. benthamiana* contains two *NbPDS* genes, Niben101Scf01283g02002.1 (*NbPDS1*) and Niben101Scf14708g00023.1 (*NbPDS2*), and the sgRNA matched *NtPDS1* but not *NtPDS2* (one bp mismatch). The *NtPDS2* genes in the regenerated plants were analyzed, revealing no off-target mutagenesis or DD insertion. To explore the off-target DD insertion that occurred in the presence of DSBs without homologous sequences with the target site, we performed whole-genome sequencing of three types of plants, in which the regenerated plant *NbPDS1* gene was (1) the same as the WT, (2) a heterozygous mutant (KO) but without TI, or (3) a bi-allelic TI (TI; Figure 1). The ssODN insertion did not occur in the WT or KO genome. In the genome of the TI regenerated plants, the DD insertion occurred not only in the DSB position created by the Cas9-gRNA RNP (Figure S5A), but also in Niben101Scf00150 (Figure S5B) and Niben101Scf06966 (Figure S5C). Using PCR, the Niben101Scf06966 and Niben101Scf00150 DNA fragments were amplified. Only Exp. 1#1 regenerated plant contained the extra TI DNA in these regions, it was not found in the other TI (#10) or KO (#2 and #6) regenerated plants (Figure S5D). Using T/A cloning for Niben101Scf00150 PCR products (Figure S5E) and Poly Peak Parser^38^ for Niben101Scf06966 (Figure S5F), these TI sequences were found to be identical to the TI sequences from whole-genome sequencing results. We genotyped Exp. 1#1 regenerated plant T_1_ seedlings and determined these two off-target TI alleles were inherited. In rice that had been transformed using Cas9-gRNA expressed from plasmid DNA via the biolistic method, quantitative PCR revealed multiple copies (2– 10 per plant) of the donor inserts in T_1_ plants, which suggested that frequent off-target DD insertion occurred in addition to the intended target site insertions.^13^ The *BoGA4* sgRNA matched *BoGA4.a* but not *BoGA4.b* (two bp mismatch). Not only targeted mutagenesis, but also off-target DD insertion occurred in *BoGA4.b* in RCBO (Figure 4C). These results indicated that the off-target DD insertion was caused by RNP and unexpected DSBs that provided an opportunity for DD insertion.

## Conclusion

In this study, we used protoplast regeneration, RNP, and ssODN to establish a simple and inexpensive DNA TI method for plant genome editing that can be used in *N. benthamiana* and RCBO by PEG-Ca^2+^-mediated protoplast transfection. In stable transformation systems, the expression of Cas protein is important for knock-in,^3^ but in this study, we used the Cas9-gRNA RNP to enhance expression together with high DD concentration to increase TI efficiency. This insertion method should be applicable to any gene target site in the genome of any plant species that can be regenerated from protoplasts without the need for antibiotic selection or phenotypic screening. Although the efficiency was low, we still obtained precise insertion regenerated plants. In the future, we will use Tandem-Repeat HDR^13^ and other methods to improve the precision and efficiency of TI and understand the precise mechanisms using ssODN.

## Acknowledgements

We thank Yu-Jung Cheng for tissue culture. We thank Yu-Lin Wu for the Illumina whole genome sequencing library preparation. We thank Miranda Loney and Plant Editors for editing. We thank the Academia Sinica Advanced Optics Microscope Core Facility for microscope imaging technical support.

## Author contributions

CSL, YCL, SL, MCS, and JS conceived and designed the experiments. CTH, QWC, and YHY performed the CRISPR-Cas9 experiments. SL performed SpCas9 protein purification. CTH, QWC, YHY, and CSL conducted the protoplast regeneration. CTH, QWC, YHY, and FHW performed the molecular biology experiments and targeted mutagenesis analysis. YCL performed whole genome sequencing and bioinformatics analysis. YCL, MCS, JS, and CSL wrote the manuscript with input from all co-authors. All authors read and approved the final manuscript.

## Author Disclosure Statement

No competing financial interests exist.

## Funding information

This research was supported by Academia Sinica, Innovative Translational Agricultural Research Administrative Office (AS-KPQ-107-ITAR-10; AS-KPQ-108-ITAR-10; AS-KPQ-109-ITAR-10), Academia Sinica Core Facility and Innovative Instrument Project (AS-CFII-108-116), and the Ministry of Science and Technology (105-2313-B-001-007-MY3; 108-2313-B-001 −011 -; 109-2313-B-001 −011 -), Taiwan.

## Supplementary Material

**Figure S1. 5-ethynyl-2′-deoxyuridine (EdU) staining of different materials.** (A) *N. benthamiana* leaves were cut into strips, placed in 1N0.3K (1/2 MS, 0.4 M mannitol, 1 mg/L NAA, 0.3 mg/L kinetin), and incubated for 0, 24, 48, or 72 hours. The incubated leaves and the protoplasts derived from these leaves were stained with EdU. *N. benthamiana* protoplasts derived from BY2 cells during cell division were used as a positive control. Bar = 50 μm. (B) Protoplasts obtained using different incubation media (1/2 MS, 0.4 M mannitol or 1N0.3K) were stained with EdU. Bar = 50 μm. (C) Protoplasts obtained using different incubation media were subjected to targeted insertion (TI) using RNP (target sequence: TTGCGATGCCTAACAAGCCAG) and donor DNA (TGCGATGCCTAACAAGCaagcttCAGGGGAGTTCAGCCGC). *NbPDS1* was subjected to PCR and analyzed by digestion with *Bst*NI (for Edited) and *Hind*III (for TI). M: DNA marker. %: the efficiency of Edited (black) and TI (red).

**Figure S2. Effect of homologous arm length and total length of the single-stranded oligonucleotide DNA donor DNA on targeted insertion in *NbPDS1* in the *N. benthamiana* regenerated plants.** (A) Protoplasts transfected with 40-nt single-stranded oligonucleotide DNA (ssODN, TGCGATGCCTAACAAGCaagcttCAGGGGAGTTCAGCCGC) and RNP (target sequence: TTGCGATGCCTAACAAGCCAG), ssODN only, RNP only, or only polyethylene glycol-Ca2+–mediated transfection. Protoplast genotypes were determined. PCR of the target gene *NbPDS1* was performed, and the PCR product was analyzed by digestion with *Bst*NI (for Edited) and *Hind*III (for TI). (B) Sequences of donor DNA (DD). The protospacer adjacent motif (PAM) is underlined. Lower case: restriction enzyme site. (C) Protoplasts transfected with different lengths of ssODN. Regenerated plant genotypes were determined. PCR of the target gene *NbPDS1* was performed, and the PCR product was analyzed by digestion with *Bst*NI (for Edited) and *Hind*III (for TI). (D) Editing and TI efficiency for different lengths of ssODN. T: total length. I: insertion. H: homologous arm. L: left arm. R: right arm. R. E.: restriction enzyme. (E) Results of Sanger sequencing of +27#6, the precise *Hind*III insertion regenerated plant (blue background). (F) BLASTN results of +27#6 *NbPDS1* (Query) and wild-type *NbPDS1* (Sbjct). Red box: DD. Yellow box: target site. Green box: inserted *Hind*III site. Black box: protospacer adjacent motif (PAM).

Figure S3. Editing efficiencies of regenerated plants from L1+L2, L1+E, E+L2 assessed using restriction enzyme digestion and ribonucleoprotein.

**Figure S4. Progeny of targeted insertion regenerated plants in *N. benthamiana*.** (A) DNA was isolated from 21 progeny seedlings from Exp. 3#7 and the target gene *NbPDS1* was amplified by PCR. The products were digested using two restriction enzymes (*Nhe*I and *Bam*HI), for which sites were added in the donor single-stranded oligonucleotide DNA (ssODN). WT: wild type PCR product digested by restriction enzyme. -: wild type PCR product without digestion. (B) Summary of the five targeted insertion (TI) regenerated plants in the T_1_ progeny. The genotype was determined as described in (A). (C) Summary of sequences of the TI alleles. Detailed sequence information for each regenerated plant is shown in Table S4 (Exp.3#7) and Table S6 (L1+E#2, 6, 9, and 10).

**Figure S5. Validation of donor DNA insertion sites by whole genome sequencing in targeted insertion regenerated plant (Exp. 1#1).** (A) An expected donor DNA insertion site of the *NbPDS1* gene on Niben101Scf01283 (red box) when aligning WGS data on the reference genome sequence. Unexpected donor DNA insertion sites on Niben101Scf00150 (B) and Niben101Scf06966 (C), respectively (black boxes). (D) The PCR products of Niben101Scf00150 (top) and Niben101Scf06966 (bottom). WT: wild type. TI: targeted insertion. KO: knock out. #: transfected regenerated plants. Letters in black: the regenerated plants for whole genome sequencing. Grey: The regenerated plants in the same transfection. (E) The Sanger sequencing results of the T/A clone contained Niben101Scf00150 PCR product of Exp. 1#1. Black box: unexpected donor DNA insertion. (F) The Sanger sequence of Niben101Scf06966 PCR product of #1 which was analyzed by Poly Peak Parser (http://yosttools.genetics.utah.edu/PolyPeakParser/)^38^. Black box: unexpected donor DNA insertion.

Table S1. Primer information.

**Table S2. Exp. 1 sequences.** (A) Summary of the TI regenerated plants. Underlined: PAM. +number: insertion length. L: left homologous arm; I: insertion; R: right homologous arm. + in Orientation column: forward insertion; -: reverse. (B) Insertion sequences of TI regenerated plants. Underlined: PAM.

**Table S3. Exp. 2 sequences.** (A) Summary of the TI regenerated plants. Underlined: PAM. +number: insertion length. –number: deletion length. L: left homologous arm; I: insertion; R: right homologous arm. + in Orientation column: forward insertion; -: reverse. (B) Insertion sequences of TI regenerated plants. Underlined: PAM.

**Table S4. Exp. 3 sequences.** (A) Summary of the TI regenerated plants. Underlined: PAM. +number: insertion length. –number: deletion length. L: left homologous arm; I: insertion; R: right homologous arm. + in Orientation column: forward insertion; -: reverse. (B) Insertion sequences of TI regenerated plants. Underlined: PAM.

**Table S5. L1+L2 sequences.** (A) Summary of the TI regenerated plants. Underlined: PAM. +number: insertion length. –number: deletion length. + in Orientation column: forward insertion; -: reverse. (B) Insertion sequences of TI regenerated plants. Underlined: PAM.

**Table S6. L1+E sequences.** (A) Summary of the TI regenerated plants. Underlined: PAM. –number: deletion length. +number: insertion length. + in Orientation column: forward insertion; -: reverse. (B) Insertion sequences of TI regenerated plants. Underlined: PAM.

**Table S7. E+L2 sequences.** (A) Summary of the TI regenerated plants. Underlined: PAM. –number: deletion length. +number: insertion length. + in Orientation column: forward insertion; -: reverse. (B) Insertion sequences of TI regenerated plants. Underlined: PAM.

**Table S8. *BoSnRK1* sequences.** (A) Summary of the TI regenerated plants. Underlined: PAM. +number: insertion length. L: left homologous arm; I: insertion; R: right homologous arm. + in Orientation column: forward insertion; -: reverse. (B) Insertion sequences of TI regenerated plants. Underlined: PAM.

**Table S9. *BoGA4.a* sequences.** (A) Summary of the TI regenerated plants. Underlined: PAM. +number: insertion length. L: left homologous arm; I: insertion; R: right homologous arm. + in Orientation column: forward insertion; -: reverse. (B) Insertion sequences of TI regenerated plants. Underlined: PAM.

**Table S10. *BoGA4.a* sequences of the RCBO TI regenerated plants by using *N. benthamiana NbPDS1* donor DNA.** (A) Summary of the TI regenerated plants. Underlined: PAM. +number: insertion length. –number: deletion length. L: left homologous arm; I: insertion; R: right homologous arm. + in Orientation column: forward insertion; -: reverse. (B) Insertion sequences of TI regenerated plants. Underlined: PAM.

## Notes

### Competing Interest Statement

The authors have declared no competing interest.

